# Protein structural context of cancer mutations reveals molecular mechanisms and identifies novel candidate driver genes

**DOI:** 10.1101/2024.03.21.586131

**Authors:** Diego Chillón Pino, Mihaly Badonyi, Colin A. Semple, Joseph A. Marsh

**Affiliations:** MRC Human Genetics Unit, Institute of Genetics and Cancer, University of Edinburgh, Edinburgh, UK

## Abstract

Advances in structure determination and computational modelling are enabling us to study the protein structural context of human genetic variants at an unprecedented scale. Here, we investigate millions of human cancer-associated missense mutations in terms of their structural locations and predicted perturbative effects. We find that, while cancer-driving mutations have properties similar to other known disease-causing mutations, this is obscured by the abundance of passenger mutations in cancer sequencing datasets. Nevertheless, by considering the collective properties of mutations at the level of individual proteins, we identify distinct mutational signatures associated with tumour suppressors and oncogenes. Tumour suppressors are enriched in structurally damaging mutations, consistent with loss-of-function mechanisms. In contrast, oncogene mutations tend to be structurally mild, reflecting selection for gain-of-function driver mutations and against loss-of-function mutations. Although oncogenes are difficult to distinguish from genes with no role in cancer using only structural damage, we find that an alternate metric based on the clustering of mutations in three-dimensional space is highly predictive of oncogenes, particularly when mutation recurrence is considered. These observations allow us to identify novel candidate driver genes and speculate about their molecular roles, which we expect to have general utility in the analysis of cancer sequencing data.

## Introduction

Tumour progression involves a complex process of genetic mutations accumulating over time, each potentially tipping the balance towards unchecked cell growth and the evasion of the body’s defence mechanisms. The proliferation of high-throughput sequencing platforms has revolutionised our ability to detect and catalogue these genetic alterations, generating abundant mutation data in large-scale tumour profiling projects such as The Cancer Genome Atlas (TCGA)^1^, the International Cancer Genome Consortium (ICGC)^2^, and the Pan-Cancer Analysis of Whole Genomes (PCAWG)^3^, and available in databases such as the Catalogue Of Somatic Mutations In Cancer (COSMIC)^4^. Despite this wealth of data, deciphering the functional consequences of these mutations remains a formidable challenge. The ability to translate these vast amounts of genetic data into actionable insights about cancer’s molecular mechanisms and potential vulnerabilities is critical for advancing the field of oncology.

Wide variations in the numbers and types of mutations are observed across different tumours. Most of these are thought to be passenger mutations that do not significantly impact tumour growth. However, a much smaller subset of mutations, known as driver mutations, play crucial roles in tumourigenesis and are selected for during tumour progression^5,6^. In fact, most tumours are believed to possess just two to eight driver mutations^7^. Identifying these driver mutations amidst the chaotic landscape of genomic changes in tumours has been a pivotal challenge in cancer genomics, and is crucial for understanding cancer mechanisms and for the development of targeted therapies^8^. While some cancer drivers are characterized by significant structural alterations, such as chromosomal rearrangements, deletions or duplications^9^, many others result from small modifications within protein-coding regions. In particular, missense mutations, *i.e.* single nucleotide changes that results in a single amino acid substitution at the protein level, have often been shown to play crucial roles in driving cancer^10,11^. It can be difficult to distinguish drivers from passengers when considering missense mutations due to their often-subtle protein-level effects, thus motivating considerable effort to develop computational predictive methods^12–16^.

Genes harbouring cancer-driving mutations are often divided into two main classes: oncogenes and tumour suppressor genes (TSGs). Oncogenes tend to play crucial roles in promoting cell growth and cell division. In contrast, TSGs act as a safeguard by keeping cell growth and division in check, thereby protecting the organism from neoplasia^17^. Numerous studies have shown that tumourigenesis is largely driven by mutations resulting in gain of function of oncogenes along with the loss of function of TSGs^17^. The intrinsic differences in the molecular mechanisms of mutations in these two categories of genes are evident in distinct in mutational patterns^18^, hotspots^19–21^ and patterns of selection^22^.

A powerful strategy for investigating the molecular mechanisms underlying protein mutations involves consideration of their protein structural context. Recently, we investigated the protein structural differences between pathogenic missense mutations, primarily associated with Mendelian genetic disorders, that act via gain-*vs* loss-of-function molecular mechanisms^23^. In particular, we observed that pathogenic gain-of-function mutations tend to have much milder effects on protein stability and interactions within protein complexes than recessive or haploinsufficient missense mutations associated with loss of function. Moreover, gain-of-function mutations showed a much greater tendency to cluster within three-dimensional protein structures. This suggested that the molecular mechanism underlying pathogenic mutations in a gene could potentially be predicted by considering protein structural context.

The classification of cancer genes into TSGs and oncogenes closely mirrors the terminology used in rare genetic disease, where most pathogenic mutations can be classified as being associated with loss-of-function or gain-of-function mechanisms^24,25^. Given that mutations in TSGs and oncogenes are often assumed to act via loss and gain of function, respectively, we reasoned that a similar large-scale analysis of cancer-associated mutations could provide insight into their cancer-driving mechanisms. Previous work has supported this notion. For example, a tendency has been observed for TSG mutations to be more structurally damaging than oncogene mutations across relatively small sets of cancer-associated mutations^15,20,26^. Moreover, a number of studies have investigated patterns of mutational clustering and hotspots in oncogenes and TSGs, with a greater degree of clustering for oncogenes often being observed^3,19,21,26–32^.

In this study, we have investigated the protein structural context of cancer-associated missense mutations on an unprecedented scale, taking advantage of the many protein and protein complex structures that have now been experimentally determined^33^, and the recent availability of computationally predicted structural models across the entire human proteome^34^. While collectively, cancer-associated mutations show only a small tendency to be structurally damaging, we observed striking structural differences between driver mutations in TSGs compared to oncogenes. Moreover, by considering the predicted stability effects on a per-gene level, we are able to identify many known TSGs as those that are most strongly enriched in structurally damaging mutations. In contrast, while oncogene mutations tended to be structurally mild, they showed strong clustering within three-dimensional protein structures. Finally, we use both structural perturbation and our clustering metric to identify genes that exhibit the characteristic properties of TSGs and oncogenes. Overall, we show that consideration of protein structure can provide new insights into the molecular mechanisms underlying cancer-associated mutations, and can potentially identify novel cancer-driving genes.

## Results and Discussion

### Cancer-associated missense mutations are enriched for structurally damaging variants

To investigate the protein structural context of cancer-associated mutations, we first downloaded all missense mutations from the Cancer Mutation Census (CMC)^35^, an ongoing project branching from the COSMIC project^4^, which we refer to as the “*cancer-all”* set of mutations in this study. These are somatic mutations that have been identified in tumour samples, but are not necessarily important for tumourigenesis; we expect many of these to be passenger mutations. We also considered a much smaller subset of the CMC comprising only those annotated for their relevance in cancer within the Cancer Gene Census (CGC), which we refer to as the “*cancer-driver”* set^35^. For comparison, we included missense mutations classified as pathogenic and likely pathogenic in ClinVar^36^ across all human protein-coding genes as the “*pathogenic”* set, and non-pathogenic missense variants observed in the human population from gnomAD v2.1^37^ as the “*putatively benign”* set, as done previously^23,38^. Next, we mapped missense mutations from the four groups to experimentally determined protein structures from the Protein Data Bank (PDB)^33^ to AlphaFold2 predicted models^34^. The total numbers of mutations in each group are provided in **Table 1**.

**Table 1.** Number of protein-coding genes and missense mutations present in the different datasets used in this study, considering those present in PDB structure or AlphaFold models.

| Dataset | PDB structures |  | AlphaFold models |  |
| --- | --- | --- | --- | --- |
|  | Genes | Mutations | Genes | Mutations |
| *Putatively benign* | 9,029 | 1,161,381 | 19,171 | 5,607,699 |
| *Cancer-all* | 8,402 | 643,008 | 17,905 | 2,675,629 |
| *Cancer-driver* | 624 | 2,080 | 1,087 | 3,348 |
| *Pathogenic* | 2,198 | 30,271 | 3,940 | 47,697 |

In **Fig. 1A**, we investigate the locations of missense mutations from the different groups within PDB structures, classifying each mutation on the basis of whether it occurs in the protein interior, surface or at an intermolecular interface. It is well known that pathogenic mutations tend to be enriched at protein interior and interface residues, as mutations at these positions are more likely to be disruptive to protein structure^39,40^. This is confirmed here, with 80% of the pathogenic set of mutations occurring at interior and interface positions, compared to only 55% of the *putatively benign* mutations. Interestingly, the *cancer-all* mutations are very similar to the *putatively benign* mutations in distribution, with only a small, albeit highly significant, enrichment at interior and interface positions (57%, p = 1.63 × 10^-97^, Fisher’s exact test). In contrast, the *cancer-driver* mutations are intermediate, with 73% occurring at interior and interface residues. Interestingly, the *cancer-driver* group is slightly enriched in interface mutations (33%) compared to the pathogenic group (30%, p = 5.58 × 10^-4^). This is consistent with previous work demonstrating enrichment of cancer-associated mutations at specific protein interfaces^41–43^. Similar patterns are observed when using AlphaFold models (**Fig. S1A**), although for these, we can only classify interior and surface positions.

**Figure 1.**
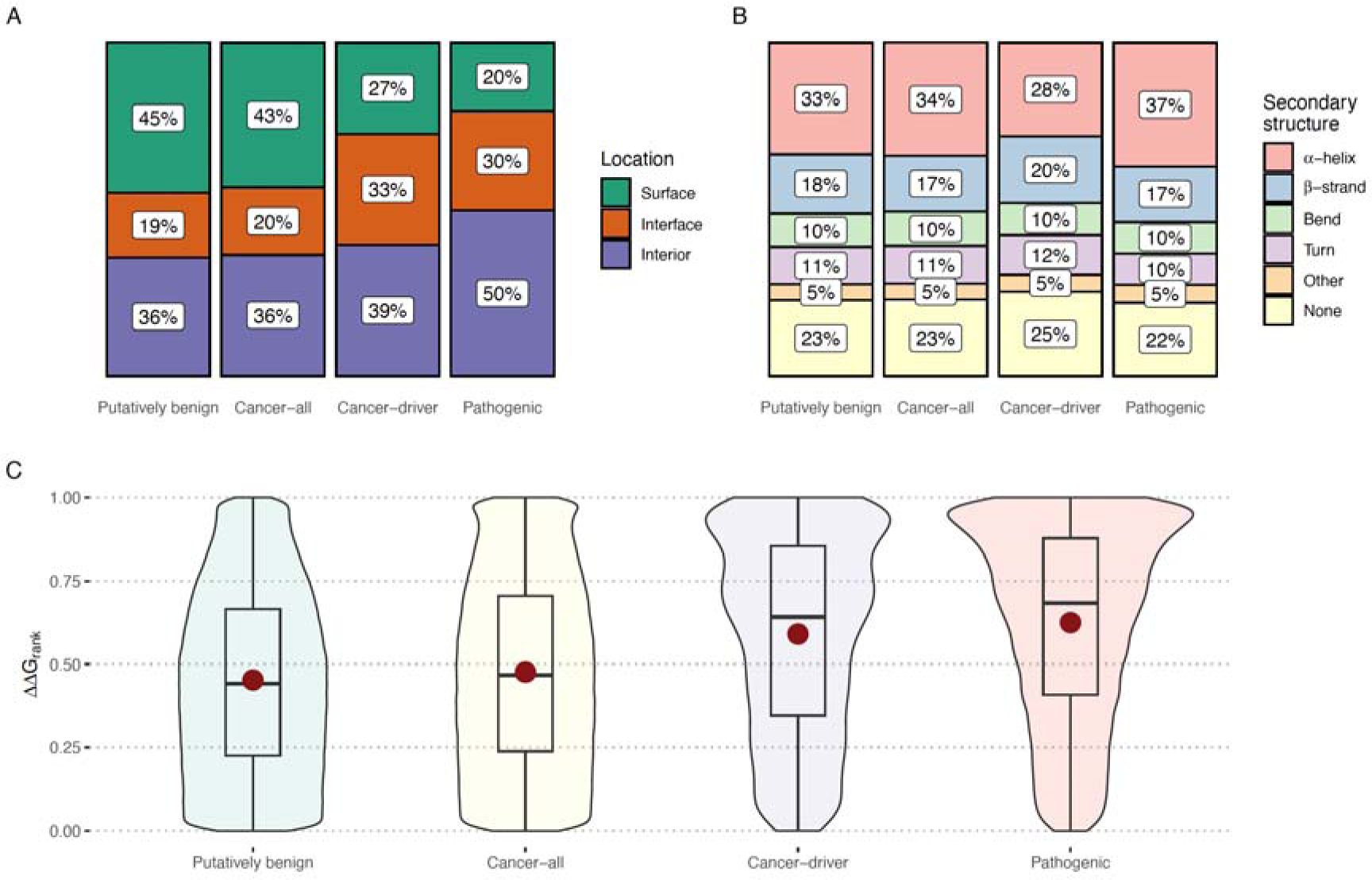
Protein structural properties of different classes of missense mutations. Putatively benign mutations are those observed in the human population (gnomAD) without a reported disease association. *Cancer-all* are all cancer-associated mutations from the CMC. *Cancer-driver* mutations are the subset of cancer-associated mutations annotated for their direct role in cancer. Pathogenic mutations are those annotated as pathogenic or likely pathogenic in ClinVar. **(A)** Locations of mutations within protein structures present in the Protein Data Bank (PDB), split into surface, interface and interior positions, as defined previously*^74^*. **(B)** Occurrence of mutations within different types of secondary structures. **(C)** Violin plot distributions of predicted structurally damaging effects, as measured by the ΔΔG_rank_ metric, whereby 0 represents the mildest possible single amino acid substitution in a protein, 1 represents the most damaging, and random mutations would be expected to have a mean of 0.5. The mean value of each distribution is represented with a red dot for the datasets. All comparisons between group pairs proved to be highly significantly different, with p-values < 2.2 x 10^-10^ according to Wilcoxon tests. Equivalent analyses based on AlphaFold models are shown in **Fig. S1**.

We also considered the distribution of mutations across secondary structure types within PDB protein structures (**Fig. 1B**), given the observation that α-helices and β-strands have different mutational sensitivities^44^. Interestingly, while the pathogenic mutations are significantly enriched at α-helices (37%) compared to the *putatively benign* (33%, p = 2.96 × 10^-40^, Fisher’s exact test) and *cancer-all* groups (34%, p = 2.48×10^-30^), the *cancer-driver* set is relatively deficient, with 28% of mutations occurring at α-helical positions (p = 3.34 × 10^-10^ *vs cancer-all*). Moreover, while all other mutation groups are nearly identical across other secondary structure classes, the *cancer-driver* group is enriched at β-strand positions (20% *vs* 17% for *cancer-all*, p = 1.5 × 10^-5^), and at regions without regular structure (25% *vs* 23% for *cancer-all*, p = 3.53 × 10^-3^). Similar trends are also seen for the AlphaFold models (**Fig. S1B**), although the *cancer-driver* group is deficient in mutations at positions with regular structure compared to the *cancer-all* group, which we speculate may be related to the treatment of disordered regions by AlphaFold. Overall, it appears that, while the differences in secondary structure between the mutation groups are fairly modest, the *cancer-driver* mutations do appear to have some distinct properties relative to both the *pathogenic* and *cancer-all* groups.

Finally, we modelled the effects on protein stability by using FoldX^45^ to calculate ΔΔG values for all mutations. Previous work has demonstrated that FoldX outperforms other stability predictors in the identification of disease mutations^46^ and shows higher correlations with deep mutational scanning data^47^, providing insights into the molecular mechanisms underlying pathogenic and cancer-associated mutations^23,28^. Because FoldX tends to occasionally output extreme outlier ΔΔG values, and because different proteins can have different intrinsic propensities for destabilising mutations, we introduced a rank normalised metric we call ΔΔG_rank_ for easier visualisation and comparison between proteins. First, for a given human protein, we use FoldX to calculate ΔΔG values for all possible missense mutations (*i.e.* all single amino acid substitutions possible via single nucleotide changes). These are then sorted based on absolute ΔΔG values, as these have been found to show slightly stronger correspondence with disease than raw ΔΔG^46^. Absolute ΔΔG values are then normalised from 0 to 1, with 0 representing the mildest possible missense mutation for a protein in terms of its effect on protein stability, and 1 representing the most structurally damaging. This scale benefits from being highly interpretable, with a mean value of exactly 0.5 across all possible mutations. Thus, in the absence of any selection, an average ΔΔG_rank_ value of around 0.5 would be expected for a set of random mutations. For the PDB structures, we compute the ΔΔG_rank_ using full complex structures, when available, as the inclusion of intermolecular interactions considerably improves the explanatory value of ΔΔG values^23,47^. For AlphaFold models, we can only evaluate the impact of missense mutations in protein monomers.

In **Fig. 1C**, we compare the distributions of ΔΔG_rank_ values calculated from PDB structures for our different mutation datasets. Consistent with previous observations, *pathogenic* mutations are significantly more structurally disruptive (mean ΔΔG_rank_ of 0.62) than *putatively benign* mutations (mean ΔΔG_rank_ of 0.45). The mean ΔΔG_rank_ value for the *putatively benign* mutations of <0.5 is consistent with some level of purifying selection for variants observed in the human population, whereas the mean ΔΔG_rank_ >0.5 for the *pathogenic* mutations implies that they are more structurally damaging than would be expected by chance. Interestingly, the *cancer-all* mutations are overall very similar to what would be expected for random missense changes, with a mean ΔΔG_rank_ of 0.48. In contrast, mutations from the *cancer-driver* set have a mean ΔΔG_rank_ of 0.59, suggesting that they are enriched in structurally damaging mutations, but are overall significantly milder than the pathogenic set. A very similar pattern is observed using the AlphaFold models, with the *cancer-all* set having properties very close to random mutations, and the *cancer-driver* set showing a significant enrichment in protein structural damage (**Fig. S1C**).

### Tumour suppressors proteins show distinct patterns of structural damage compared to oncogenes

Our initial results show that, overall, cancer-associated missense mutations are only very slightly more damaging at a protein structural level than putatively benign mutations from gnomAD observed in the human population, consistent with the idea that the mutational landscape of tumours is dominated by passenger mutations. Interestingly, however, even those mutations with evidence for being cancer drivers are still milder than pathogenic ClinVar mutations, suggesting that, in general, cancer-driving missense mutations tend to have weaker effects on protein structure than mutations that cause Mendelian genetic disorders. Our previous work demonstrated that missense mutations that cause disease via gain-of-function mechanisms tend to induce much smaller perturbations in protein stability than those that act via a loss of function^23^. Therefore, we hypothesised that the milder protein structural effects of cancer-driving mutations are due to a greater tendency to be associated with gain-of-function effects.

First, we compare the *cancer-all* mutations from oncogenes and TSGs (**Fig. 2A**). We observe some differences in the location distributions between the two groups, with TSGs being moderately enriched in mutations at interior positions (40% *vs* 35%, p = 3.5 × 10^-31^ Fisher’s exact test) and slightly enriched at interface positions (20% *vs* 19%, p = 3.69 × 10^-3^). Moreover, there is a highly significant tendency for TSG mutations to be more structurally damaging than oncogene mutations, as measured by ΔΔG_rank_ values, although the overall difference between the two groups is small, with a mean of 0.48 for the oncogenes and 0.50 for the TSGs (**Fig. 2B**). Similar trends are observed for AlphaFold models (**Fig. S2**).

**Figure 2.**
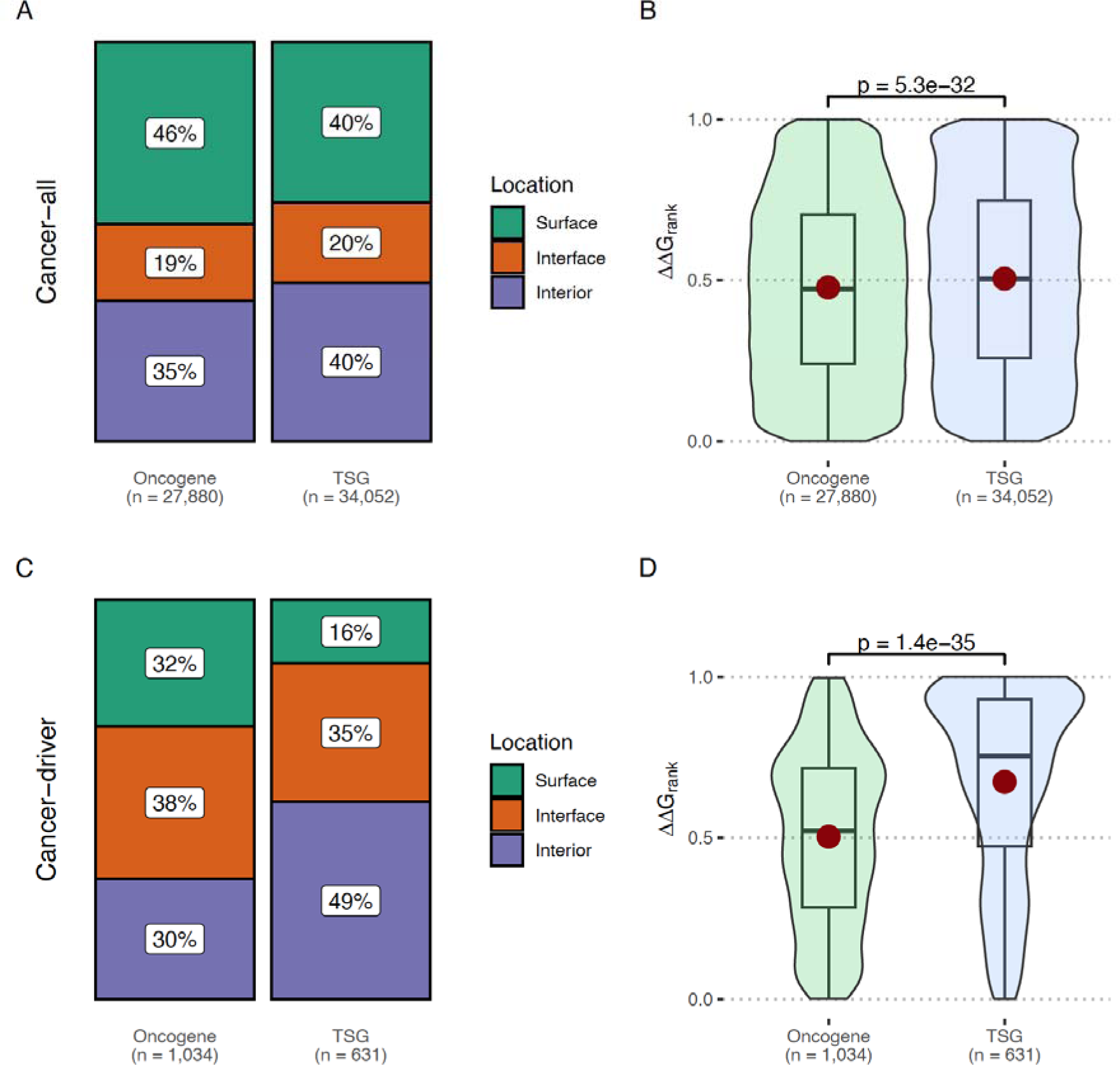
Protein structural properties of cancer-associated mutations in oncogenes and tumour suppressors. **(A)** Locations of all cancer-associated mutations within PDB structures (*cancer-all* dataset) and **(B)** distributions of predicted structural damage, as represented by ΔΔG_rank_ values. **(C)** Locations of cancer-associated mutations with an annotated role in cancer (*cancer-driver* dataset) and **(D)** distributions of ΔΔG_rank_ values. *P*-values are calculated using Wilcoxon tests. Equivalent analyses based on AlphaFold models are shown in **Fig. S2**.

Next, we consider the *cancer-driver* mutations **(Fig. 2C)**. Both groups are highly represented at interface locations, which account for 38% and 35% of the mutations in oncogenes and TSGs, respectively. TSGs also have far more mutations at interior positions (49%) compared to oncogenes (30%, p = 5.65 × 10^-5^, Fisher’s exact test). When considering ΔΔG_rank_, the difference between the two groups is striking, with a mean of 0.66 for TSGs *vs* 0.47 for oncogenes (**Fig. 2D**). Thus, the cancer-driving missense mutations in TSGs tend to be even more damaging than the pathogenic mutations. This supports the idea that the structurally milder nature of the cancer-driving mutations, when considered collectively, is a consequence of their lower tendency to be associated with loss-of-function molecular mechanisms, due to the oncogenic nature of many cancer-driving mutations. In other words, the balance between loss-of-function *vs* gain-of-function effects appears to be shifted towards gain of function for cancer-driving mutations compared to pathogenic mutations associated with genetic disease.

Although the differences in structural damage between oncogenes and TSGs were minimal in the *cancer-all* dataset when considering these groups collectively, we wondered if we could identify specific protein-coding genes enriched in structurally damaging or structurally mild mutations. For each protein, we calculated the difference between the mean ΔΔG_rank_ for the *cancer-all* mutations, and the mean ΔΔG_rank_ for all other possible (but not observed) missense mutations. Proteins with a ΔΔG_rank_ difference greater than 0 are relatively enriched in mutations that are structurally damaging compared to what would be expected if mutations occurred randomly without selection, whereas those with a negative ΔΔG_rank_ difference will have mutations that are less structurally disruptive than would be expected. We show ΔΔG_rank_ difference values across all proteins based on PDB structures (**Fig. 3A**) and AlphaFold models (**Fig. 3B**) as a volcano plot, where the Wilcoxon *p-*value represents the significance of the difference between observed and unobserved mutations.

**Figure 3.**
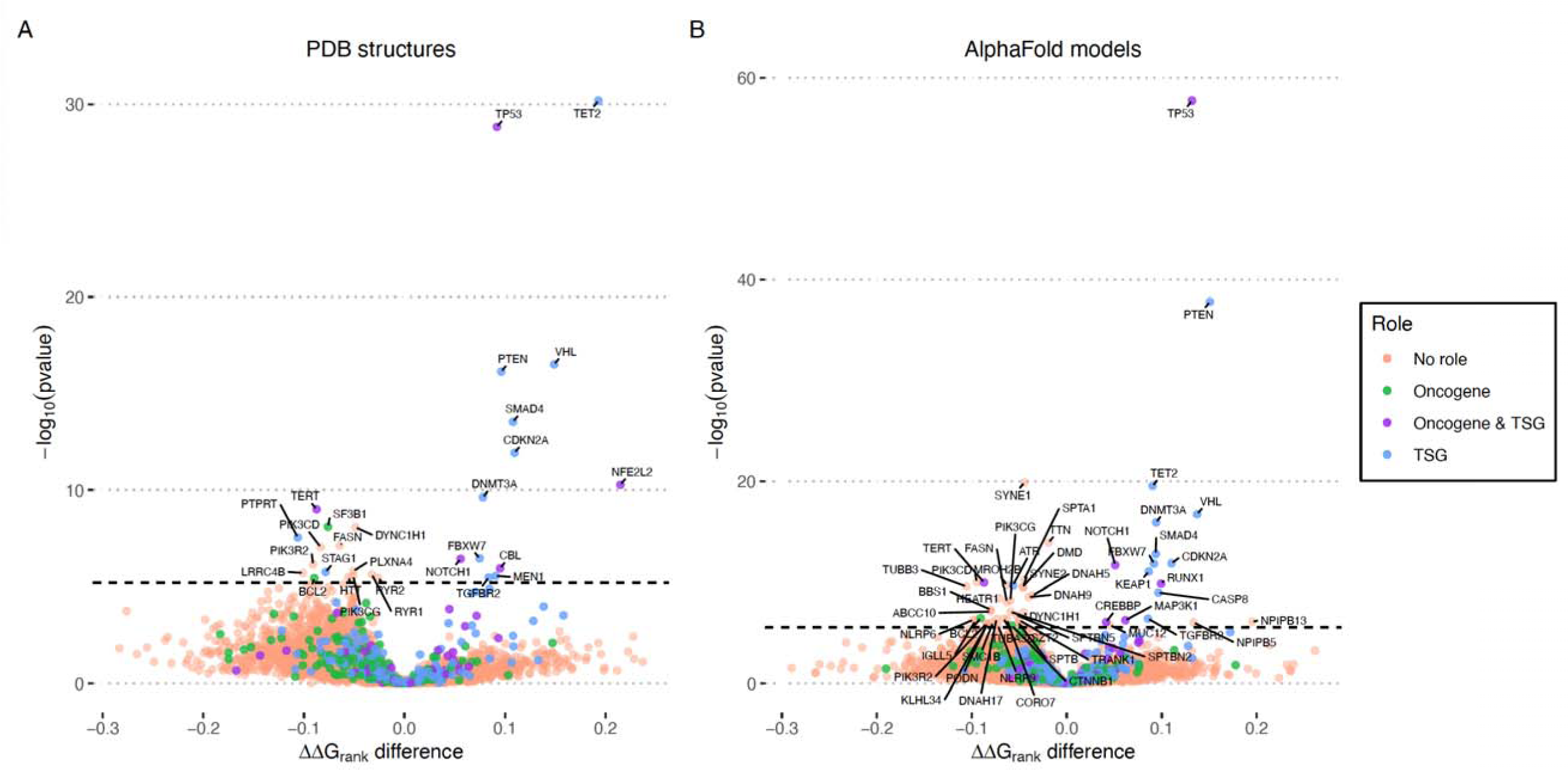
Gene-level enrichment in structurally damaging and structurally mild cancer-associated missense mutations. For each of the 8402 human protein-coding genes with a PDB structure **(A)** or 17905 with an AlphaFold model **(B)**, we plot the difference between the mean ΔΔG_rank_ for mutations observed in the *cancer-all* dataset and for other possible missense mutations not present in the dataset. Proteins with positive ΔΔG_rank_ difference values are enriched in structurally damaging mutations, in that the average of the observed cancer-associated mutations is more destabilising than the average of the possible but unobserved mutations. In contrast, proteins with negative ΔΔG_rank_ difference values are enriched in structurally mild mutations, in that the average of the observed mutations is less destabilising. The *p*-value of this difference is calculated with the Wilcoxon test. The horizontal dashed bar represents the threshold for a statistically significant values, as result of a Bonferroni correction: 6.26×10^-6^ for the PDB structures and 2.88×10^-6^ for the AlphaFold models. Proteins are coloured based on their classified role in the Cancer Gene Census (CGC).

Proteins on the right sides of the volcano plots are enriched in structurally damaging mutations. For those with statistically significant *p*-values (above the dashed line), this implies that there has been selection for damaging mutations. In other words, structurally damaging mutations in these proteins are expected to drive cancer. Remarkably, all of the proteins with the most significant enrichments in structurally damaging mutations, using both PDB structures and AlphaFold models, have known tumour suppressor activity, including TET2, TP53, VHL, PTEN, SMAD4, CDKN2A, NFE2L2 and DNMT3A. Even below the strict statistical significance threshold (p < 6.26 × 10^-6^), which accounts for multiple testing, there is a notable enrichment of known TSGs on the right side of the plot, suggesting that this approach could be useful for identifying genes with putative tumour suppressor functions.

Although the most significantly enriched proteins occur on the right side of the volcano plots, reflecting strong selection for structurally damaging mutations in certain proteins, there are far more proteins with negative ΔΔG_rank_ difference values. There are two potential explanations for these proteins, which are enriched in structurally mild mutations. First, to some extent, this is likely to reflect positive selection for cancer-driving mutations that are not structurally damaging, *e.g.* gain-of-function mutations. Indeed, the two most significantly enriched proteins in the PDB analysis, TERT and SF3B1, have known oncogenic activity. However, purifying selection against structural damage mutations may be an even greater contributor to the enrichment in structurally mild mutations. In cancer-driving oncogenes, damaging mutations that cause a loss of function are likely to be strongly selected against. In addition, proteins that have no specific role in cancer, but which are important for cellular growth or viability, are also likely to experience cancer-level selection against structurally damaging loss-of-function mutations. Thus, a statistical enrichment in structurally mild mutations does not necessarily imply an oncogenic role.

Several proteins classified as TSGs also appear on the left sides of the plots, including PTPRT, STAG1 and ATR. This suggests that the cancer-driving effects of mutations in these proteins are unrelated to loss of function induced by intramolecular destabilisation. One possible explanation is that damaging mutations in these TSGs disrupt other aspects of function, such as protein interactions. The AlphaFold analysis, being based only on monomeric models, will not account for any interaction-disrupting effects. While the PDB analysis does include many experimentally determined protein complex structures, these do not include all biologically relevant interactions. Thus, some mutations that have little apparent structural effect in our analysis may actually be highly damaging to specific protein interactions.

### Oncogene mutations show characteristic clustering in three-dimensional space

In our previous work, we introduced a novel protein structural metric, the Extent of Disease Clustering (EDC), that showed remarkably strong discrimination between genes associated with gain-of-function *vs* loss-of-function mechanisms^23,48,49^. EDC is a simple measure that quantifies the extent of disease mutation clustering within a three-dimensional protein structure. An EDC value greater than one indicates that disease mutations tend to be close to each other within the structure, while a value of one would be expected if the disease mutations were randomly distributed throughout the protein. Therefore, given the association of oncogenes and TSGs with gain-of-function and loss-of-function mutations, respectively, we wondered whether EDC values would also be useful for the identification of cancer-associated genes, and for the discrimination between oncogenes and TSGs. To illustrate, in **Fig. 4A**, we show two examples of mutation distributions within protein structures and their associated EDC values. For the oncogene KRAS, known cancer-driving missense mutations are observed to be highly clustered on the protein structure, resulting in a high EDC value of 1.69. In contrast, for the tumour suppressor SDHB, the known driver mutations are spread throughout the protein, resulting in a low EDC of 0.85.

**Figure 4.**
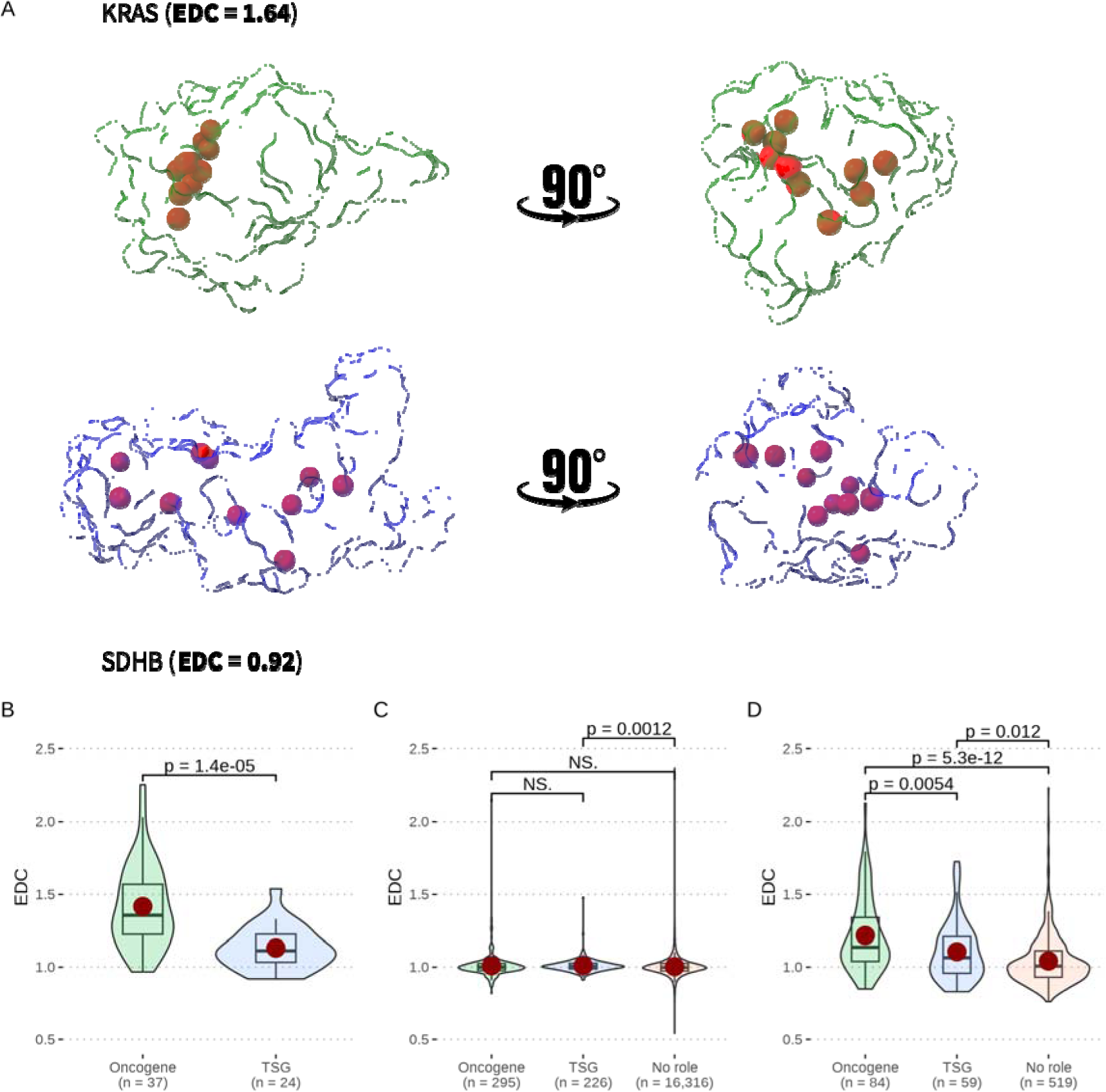
Clustering of cancer-associated mutations in three-dimensional space. **(A)** Location of *cancer-driver* mutations for an oncogene (KRAS) and TSG (SDHB), highlighting their remarkably different clustering, as reflected by the high Extent of Disease Clustering (EDC) value for KRAS, and low EDC value for SDHB. **(B)** Distribution of EDC values calculated from mutations from the *cancer-driver* dataset, split into those cancer-associated genes classified as having oncogene activity, and those genes only classified with TSG activity. **(C)** Distribution of EDC values calculated from the *cancer-all* dataset. **(D)** Distribution of recurrent EDC values calculated from the *cancer-all* dataset. *P*-values are calculated using Wilcoxon tests.

First, we considered EDC values for known cancer-driving mutations from the *cancer-driver* dataset (**Fig. 4B**), revealing a strong, highly significant tendency for the mutations in oncogenes to be more clustered than those from TSGs. We considered proteins with mutations present at five or more residues, the same threshold as we have used in recent studies^48,49^, although our results are similar across different minimum residue thresholds (**Fig. S3A**). Interestingly, the degree of clustering appears even stronger than we previously observed for pathogenic missense mutations^23^. The oncogenes had a median EDC of 1.48, compared to 1.25 previously observed for gain-of-function mutations. The TSGs had a mean EDC of 1.13, similar to the value of 1.09 observed for loss-of-function missense mutations in autosomal dominant genes.

We next calculated EDC values from the *cancer-all* dataset and thus consider the clustering properties of driver and passenger mutations collectively. We observed similar distributions of EDC values for oncogenes and TSGs, as well as for genes with no known role in cancer (**Fig. 4C**). In fact, the large majority of proteins show very little deviation from EDC values of one, which is markedly different from the pattern we observed when considering pathogenic missense mutations from ClinVar^23,48^. Thus, it appears that the large number of passenger missense mutations in this dataset likely obscures our ability to detect any signs of clustering using the EDC metric, even in known oncogenes and TSGs. The clustering of known driver mutations is of no utility for us for identifying novel candidate cancer-driving genes.

To overcome this limitation, we next limited our analysis to a subset of the *cancer-all* dataset, considering only those missense mutations that are recurrent. This allows us to include far more genes than with the *cancer-driver* dataset, many of which have no known role in cancer (**Fig. 4D**). To calculate recurrent EDC values, we considered only residues mutated at least seven times in *cancer-all* data, although our results are robust to different recurrence thresholds (**Fig. S3B**). We observe significantly higher recurrent EDC values in the in the oncogenes compared to TSGs, although the extent of clustering is somewhat less pronounced than observed for the driver mutations alone, with a mean EDC of 1.16 for the oncogenes compared to 1.07 for the TSGs. Notably, we also observe EDC values in both oncogenes and tumour suppressors to be significantly higher compared to genes with no known cancer role, suggesting that some degree of clustering does occur in TSGs, but to a less extent than in oncogenes. This may be related to damaging missense mutations being more likely in certain regions of tumour suppressors, *e.g.* around functionally important sites, and is consistent with the previous observation of clustering in other TSGs, e.g. SPOP^50^.

### Identification of novel candidate tumour suppressors and oncogenes using protein structural information

As both structural disruption and high three-dimensional clustering of missense mutations appear to be predictive of genes that have roles in cancer, we explored the potential of these properties to prioritise novel candidate TSGs and oncogenes. First, in **Fig. 5A**, we show the 50 proteins most significantly enriched in structurally damaging mutations, combining both the PDB and AlphaFold analyses. All of the 15 highest ranked proteins, and 34 out of the top 50, have known tumour suppressor activities, demonstrating the strong potential of this approach for identifying putative cancer-driving genes.

**Figure 5.**
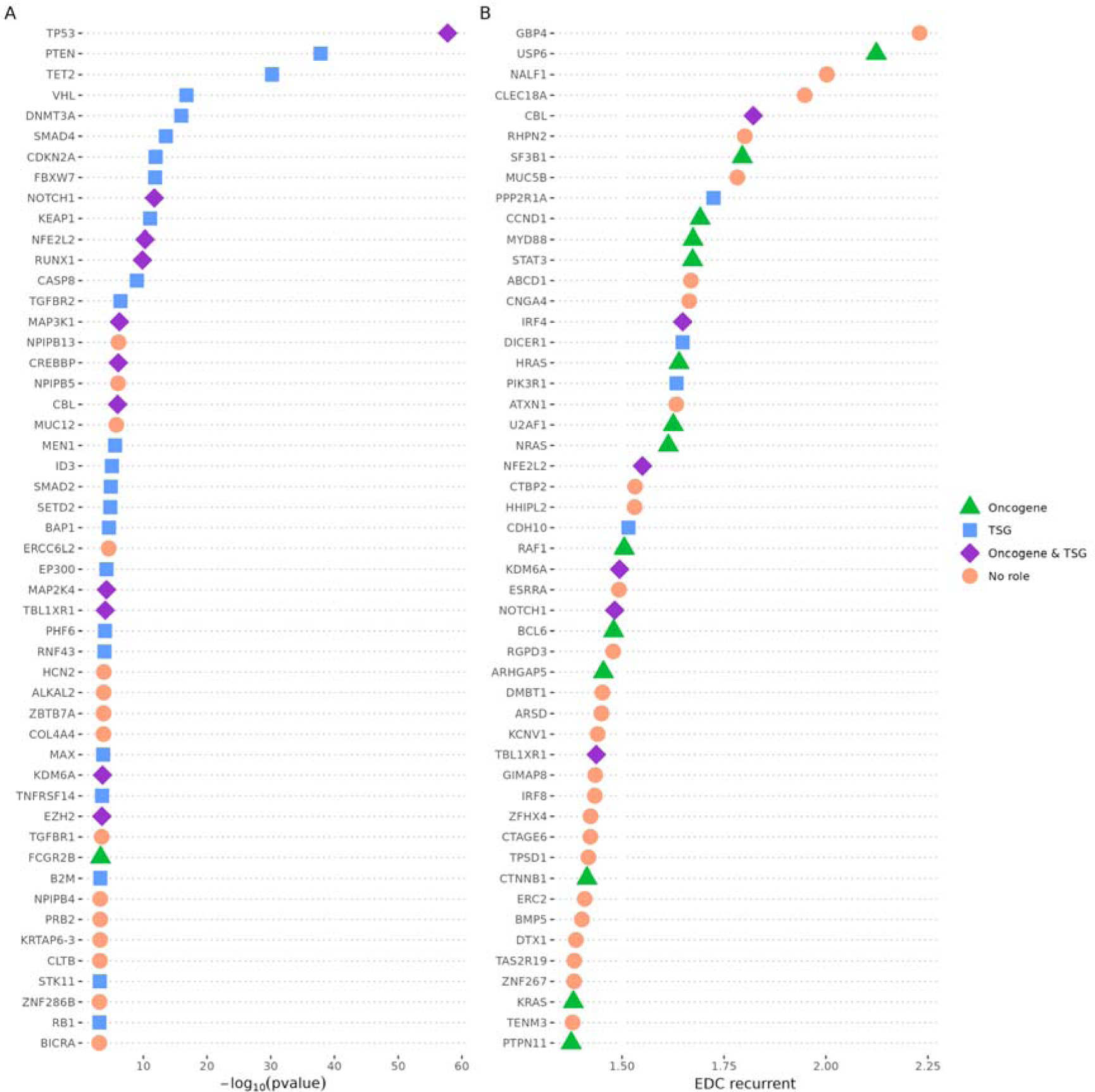
Prioritisation of putative cancer-driving genes. (A) Top 50 human protein-coding genes enriched in structural damaging mutations, *i.e.* those with positive ΔΔG_rank_ difference values. For these proteins, the observed missense mutations in the *cancer-all* dataset are significantly more destabilising than the possible but unobserved mutations. *P*-values from both the PDB and AlphaFold analyses in Fig. 3 are included, with the most significant value from either analysis selected for each protein to be used in this ranking. (B) Top 50 human protein-coding genes with the highest recurrent EDC values, based on recurrent mutations from the *cancer-all* dataset, as in Fig. 4D.

Next, we explored those proteins with no role in cancer as classified in the CGC. The most significantly enriched of these proteins, NPIPB13 and NPIPB5 (ranking 16^th^ and 18^th^ overall, respectively), are both members of the primate-specific nuclear pore complex interacting protein (NPIP) family. Very little is known about the normal biological function of this family, or their potential role in human disease. One recent study has linked NPIPB5 expression to prognosis and patient survivability in renal cell carcinoma^51^, while another found NPIPB13 expression to be weakly associated with microvascular invasion in hepatocellular carcinoma^52^. The closely related NPIPB4 is also listed in **Fig. 5A**, ranking 43^rd^ overall. While the previous limited known biological role or cancer association for these proteins may argue against a tumour suppressor function, we find it interesting that similar patterns of enrichment in damaging mutations are observed across all three of these closely related homologues, suggesting that it could be worthy of further investigation.

MUC12 was the next most significantly disrupted gene with no cancer classification, ranking 20^th^ overall. Previously, its expression was found to be significantly lower in colorectal cancer tissues, indicative of potential tumour suppressor activity^53^. In contrast, other research suggested that MUC12 was overexpressed in renal cell carcinoma^54^. Its high significance is influenced by its long length (5478 amino acids), and the effect size is relatively small, but our results suggest that there may be a tendency of structurally damaging missense mutations in this protein to drive cancer.

ERCC6L2 ranks 26^th^ overall, and appears to be involved in DNA repair processes^55^, with recessive protein null mutations being associated with bone-marrow-failure syndrome^56^. It also possesses two domains that commonly occur in known TSGs: the *Helicase_C* domain is found in nine well-established TSGs, and the *SNF2-rel_dom* domain is found in three. Notably, patients with bone-marrow-failure syndrome have also been observed to be at a high risk of developing acute myeloid leukemia^57^. Thus, ERCC6L2 seems to be a strong candidate as a putative tumour suppressor.

ZBTB7A, ranking 34^th^ overall, is a transcriptional repressor involved in cell proliferation and differentiation^58^. The protein has an N-terminal BTB dimerisation domain. BTB domains are known to be strong drivers of cotranslational assembly^59^, a process that lessens the likelihood of observing a disease mechanism other than loss of function^48^. Consistently, heterozygous variants in ZBTB7A have been linked to a neurodevelopmental phenotype and are suggested to cause loss of function^60^. While the gene has not yet been classified with a cancer role in the CGC, it is increasingly being recognised as a potential cancer driver^61^. For example, somatic loss-of-function mutations in ZBTB7A cause elevated glycolysis in human cancer^62^, and loss of one copy or the C-terminal zinc finger domains have been associated with acute myeloid leukaemia^63^. Thus, while in the case of ZBTB7A our method does not offer a completely novel target, it nevertheless further supports its role in cancer and provides validation of our approach.

We also investigated the use of recurrent EDC values to identify cancer-driving genes. In **Fig. 5B**, we show the top-50 proteins with the highest EDC values, based on recurrent mutations from the *cancer-all* datasets, as used in **Fig. 4C**. Of these, 24 are classified as known cancer-driving genes in the CDC, with 14 being oncogenes, 6 having both oncogene and TSG activity and 4 being TSGs. This is consistent with our observation that, while oncogenes show the highest degree of mutation clustering, there is also significant clustering in TSGs. Although known cancer drivers are not quite as highly enriched as for the structural damage analysis, this does appear to be a promising strategy for identifying putative cancer-driving genes.

GBP4 shows the strongest clustering among all proteins, with a recurrent EDC value of 2.23. Examination of the underlying mutational data shows a cluster of highly recurrent mutations at a small stretch from residues 541-551 in the coiled-coil domain, supportive of a potential cancer-driving activity associated with this region. While the precise role of GBP4 in cancer is still unclear, there has been some previous work suggesting its involvement^64^; in particular, it has been observed to be upregulated in certain tumour types^65^.

Another of our top hits with no cancer classification, CNGA4, has a recurrent EDC of 1.66 and features two domains, transmembrane ion transport domain (*Ion_trans*) and cyclic nucleotide-monophosphate binding domain (*cNMP_binding*) that are also found in two known oncogenes, CACNA1D and PRKAR1A, respectively. Recurrent *cancer-all* mutations in CNGA4 are limited to 5 distinct residues that exhibit high clustering in the *Ion_trans* domain. Leveraging the previously modelled tetrameric structure of CNGA4^66^, our analysis revealed that the mutations cluster at the channel pore, suggesting a potential gain-of-function effect. Since CNGA4 is important for transduction of odorant signals^67^, it is possible that mutant proteins are advantageous to chemotaxis-mediated processes in cancer^68^.

Four proteins occur in the top 50 for both enrichment in structurally damaging mutations (**Fig. 5A**) and recurrent EDC (**Fig 5B**): NOTCH1, KDM6A, CBL and TBLXR1. Interestingly, all of these are classified as both oncogenes and TSGs in the CGC. Thus, the combination of high structural damage and clustering in three-dimensional space represents a strong indicator of genes with both oncogenic and tumour suppressor activity.

As discussed earlier, enrichment in structurally mild mutations is not nearly as predictive of cancer association as enrichment in damaging mutations. In **Fig. S4**, we show the top 50 genes most significantly enriched in structurally mild mutations (*i.e.* those from the left side of the volcano plots in **Fig. 3**). Of these, only eight have a known cancer role, including four oncogenes, three TSGs, and one with both activities. Thus, we think it is unlikely that a large proportion of the remaining 42 genes with no classifications are playing important roles in cancer. Many of the proteins most significantly enriched for mild mutations are of large size. Very long proteins like SYNE1 and TTN show fairly modest negative ΔΔG_rank_ values but appear highly significant in the AlphaFold analysis (**Fig. 3B**), due to the large number of mutations. Similarly, the very large calcium channels RYR1 and RYR2 are both significantly enriched in the PDB analysis, but it seems unlikely they represent true cancer drivers. Interestingly, the beta-tubulin gene TUBB3 is also highly enriched in structurally mild mutations. Given that TUBB3 expression is strongly associated with resistance to anti-microtubule chemotherapeutics^69^, this could possibly result from selection against damaging mutation in patients who have undergone chemotherapy, or selection for mutations that confer greater resistance.

## Conclusions

This study investigates the protein structural context of cancer-associated missense mutations. A major challenge associated with this is the fact that datasets of cancer mutations are inevitably dominated by passenger mutations. Thus, although we find that properties of known cancer-driving mutations are similar to the properties of other pathogenic missense mutations, analyses of all cancer-associated mutations together reveal much weaker trends. Despite this, we can observe meaningful effects by considering the collective gene-level properties of cancer-associated mutations. In particular, mutations in tumour suppressors tend to be significantly enriched in structural damage, similar to pathogenic loss-of-function missense mutations. In contrast, mutations in oncogenes tend to be structurally mild but show strong clustering within three-dimensional protein structures, similar to gain-of-function disease mutations. By searching for genes enriched in these mutational properties, we can identify novel candidate cancer drivers and obtain insight into the molecular mechanisms by which mutations in these genes might act.

The findings here are made possible by the huge number of structural models for human proteins now available. Analyses based on PDB structures are somewhat limited due to the relatively small number of human proteins with published experimentally determined structures. Nevertheless, this PDB-level analysis has allowed us to assess the structural context of 24% of the *cancer-all* missense mutations and 62% of the *cancer-driver* mutations (**Table 1**). The key advantage of the PDB-based analyses is that they can consider the effects of intermolecular interactions, given the fact that most human proteins are able to assemble into complexes^70^. Thus, the trends observed for certain proteins were markedly higher when using the PDB structures. For example, TET2 achieves much higher significance based on the PDB structures compared to the AlphaFold models in **Fig. 3**, likely because the predicted effects of many missense mutations will be greater due to their disruptive effects on the interaction with DNA. In contrast, the AlphaFold models have the advantage of being available for all human proteins, and we note that many of the interesting hits we identified in our search for putative cancer drivers are in proteins for which experimentally determined structures are not available.

A key limitation of this study is its broad focus on pan-cancer effects, which might overlook the nuances of tissue-specific oncogenesis. While this maximises statistical power for the purposes of this study, we acknowledge that many cancer-driving genes and mutations will have a strong tissue specificity. Tumourigenesis starts as a localised process in a specific tissue, and all the subclonal populations resulting from the original retain features and characteristics of the original cell line. This results in tumours displaying traceable transcriptomes and interactomes, even after metastasis, and distinctive molecular mechanisms linked to their aetiology. For instance, approximately 30–50% of colorectal cancer tumours have a mutated *KRAS* gene^71^, whereas it has been observed to be mutated in 90% of pancreatic cancers of all grades^72^, and the mutation signatures differ between them as well. Future work should focus on the tissue specificity of the phenomena observed here, and this will be facilitated by the rapid growth in available cancer sequencing data.

A crucial focus of this study has been on predicting the effects of missense mutations on protein stability, based on our previous observation of the power of this for explaining molecular mechanisms^23^. However, computationally predicted ΔΔG values are limited in their utility for identifying pathogenic missense mutations compared to evolution-based variant effect predictors (VEPs)^46^. It is possible that applying the approach we have used here, but using state-of-the-art VEPs rather than FoldX-predicted ΔΔG values, could prove even more powerful for identifying cancer-driving genes. However, we suspect that this strategy may be less informative regarding molecular mechanisms, given that VEPs rely primarily on evolutionary information and should be relatively insensitive to whether or not functionally disruptive mutations are damaging to protein structure^23^. Moreover, given that nearly all VEPs underperform on gain-of-function compared to loss-of-function mutations^23^, and our observation that cancer-driving mutations appear to be enriched in gain-of-function effects, the ability of the current generation of VEPs to provide insight into cancer-driving mutations and genes may be limited.

Our results provide further demonstration of the utility of our EDC metric, which quantifies the extent of mutation clustering within protein structures, and has proven valuable for distinguishing between genes associated with loss-of-function, gain-of-function and dominant-negative disease mechanisms^23,48,49^. At the same time, it is interesting that no meaningful trends were observed in the *cancer-all* dataset, suggesting that this approach is highly sensitive to the noise associated with passenger mutations. While considering mutation recurrence enabled us to overcome this, it is an imperfect solution that likely loses some useful information regarding clustering. A more nuanced strategy that considers number of occurrences relative to a background mutational null model may provide a better way of identifying clustering within noisy cancer mutation datasets.

Here, we identified human protein-coding genes that we refer to as novel candidate cancer drivers. Many of these appear to be interesting, but it is very difficult to confirm any cancer-associated role they may possess. Validating these associations in independent cancer sequencing datasets will be essential for providing further confidence in our predictions. Moreover, by making our gene-level results available, including ΔΔG_rank_ difference and associated *p*-values, and recurrent EDC values, we hope that our results will guide others and provide independent evidence in the search for novel cancer-driving genes. In addition, as more cancer sequencing data becomes available, the power our approach, and the power to apply in in a tissue-specific manner, will continue to grow.

## Methods

### Data collection

All somatic missense mutations observed in tumours were retrieved from the CMC v95^35^, which was downloaded from the COSMIC portal^4^. Additional data files were also downloaded for mapping purposes from the same repository, including gene-level classification information and the CGC. Pathogenic missense mutations were retrieved from ClinVar as of 2022.10.09, including those classified as pathogenic and as likely pathogenic. Putatively benign missense variants were collected from gnomAD v2.1.1, excluding any that were present in the pathogenic set.

Only gene level classifications of “oncogene” and “TSG” were considered. Although the CMC categorises some genes as being associated with “fusion”, we ignored this given that it represents a fundamentally different type of genomic change compared to the missense mutations we are interested in. Thus, a gene classified in the CGC as “fusion” was considered as having no role in our analysis, while a gene classified as “oncogene, fusion” would be considered an oncogene.

For the *cancer-driver* dataset, we only included the subset of CMC missense mutations directly annotated as having a role in cancer. Although these mutations are assigned a tier level of 1-3 in the CMC based on strength of evidence, we grouped them all together here due to the limited size of our dataset.

### Structural dataset

The PDB analysis was performed using the same analysis pipeline as previously described^23^. In short, protein structures were downloaded from the Protein Data Bank on 2022.08.05, using the first biological assembly for each entry. All missense mutations were mapped to structures, considering regions with >90% sequence identity to the human protein over a region of at least 50 residues. In cases where a residue maps to more than one polypeptide chain, we first selected the highest resolution structures, and in the case of ties, selected the largest biological assembly. The AlphaFold analysis used AlphaFold2 version 1 models^34^, downloaded from https://alphafold.ebi.ac.uk/download on 2021.07.27. Secondary structure of each residue was classified with DSSP^73^, and interior, surface and interface residues were defined according to relative solvent accessibility (RSA)^74^. “Interior” residues have an RSA ≤ 0.3; “Interface” residues have an RSA between 0.3 and 0.5 (0.3 < RSA < 0.5); “Surface” residues have an RSA ≥ 0.5. The “Interface” category is collapsed into “Surface” for the AlphaFold dataset.

FoldX 5.0 calculations were performed using all default parameters, with three replicates per mutation, and the ‘RepairPDB’ function run in advance. Only ‘full’ ΔΔG values based on the entire biological assembly were used for PDB structures, while AlphaFold models were monomeric. For large proteins, where multiple overlapping AlphaFold models are generated, we averaged ΔΔG values over all available models for each variant. We rank normalise absolute ΔΔG values to obtain the ΔΔG_rank_ metric, whereby the mildest |ΔΔG| is defined as being equal to 0, the highest |ΔΔG| was defined as 1, and the mean of all possible amino acid substitutions in a protein was equal to 0.5. Absolute ΔΔG values were used, based on our previous observation of their slightly improved correspondence with mutation pathogenicity^46^.

The Extent of Disease Clustering (EDC) metric was calculated as previously described^23^. For PDB structures, all residues were considered. However, for AlphaFold models, residues with low-confidence structural predictions, having predicted local distance difference test (pLDDT) values less than 70, were excluded from the calculation. This is similar to our most recent studies^48,49^, as we found that, for pathogenic missense mutations, this results in much better discrimination between proteins with loss-of-function and non-loss-of-function mutations when using EDC derived from AlphaFold models. Only proteins with mutations occurring at five or more residues were considered. For the recurrent analysis, only residues observed to have been mutated at least seven times (using the COSMIC_SAMPLE_MUTATED column in the CMC dataset) were included. Our results are robust to the choice of these thresholds (**Fig. S3**).

### Data curation and analysis

All data curation, mapping and analysis was carried out using R. RStudio was used for scripting. All three datasets mentioned in the data collection section were filtered keeping only unique missense mutations to avoid potential biases and duplicates in the results. The collection of R packages from the tidyverse and the data.table package were used to smooth and speed up the running time of the code, as well as to significantly increase the legibility of the code. The furrr and future packages were used to implement parallel computing and optimise code runtime. Data visualisation was achieved using the R package ggplot2 and extensions based on it, namely ggstatsplot (for in-plot statistics), ggrepel (for non-overlapping labelling), and patchwork (for composing multi-pane plots). ChimeraX^75^ (v1.5) was additionally used to visualise variant clustering in a 3D context. Some discrepancies and inconsistent annotations resulted in dropping a very small number of mutations for each dataset (≈1% variant loss for the three databases), mostly found in fusion genes.

## Acknowledgements

We thank Benjamin Livesey and Lukas Gerasimavicius for their helpful comments on the manuscript. This project was supported by funding to JAM from the European Research Council (ERC) under the European Union’s Horizon 2020 research and innovation programme (grant agreement No. 101001169), a Lister Institute Research Prize Fellowship and by the Medical Research Council (MRC) Human Genetics Unit core grant. (MC_UU_00035/9). This work has made use of the resources provided by the Edinburgh Compute and Data Facility (ECDF) (http://www.ecdf.ed.ac.uk/).

## Data Availability

Complete datasets associated with the analyses in this study are available at https://osf.io/vk68d/

## Supplemental Figures

**Figure S1.**
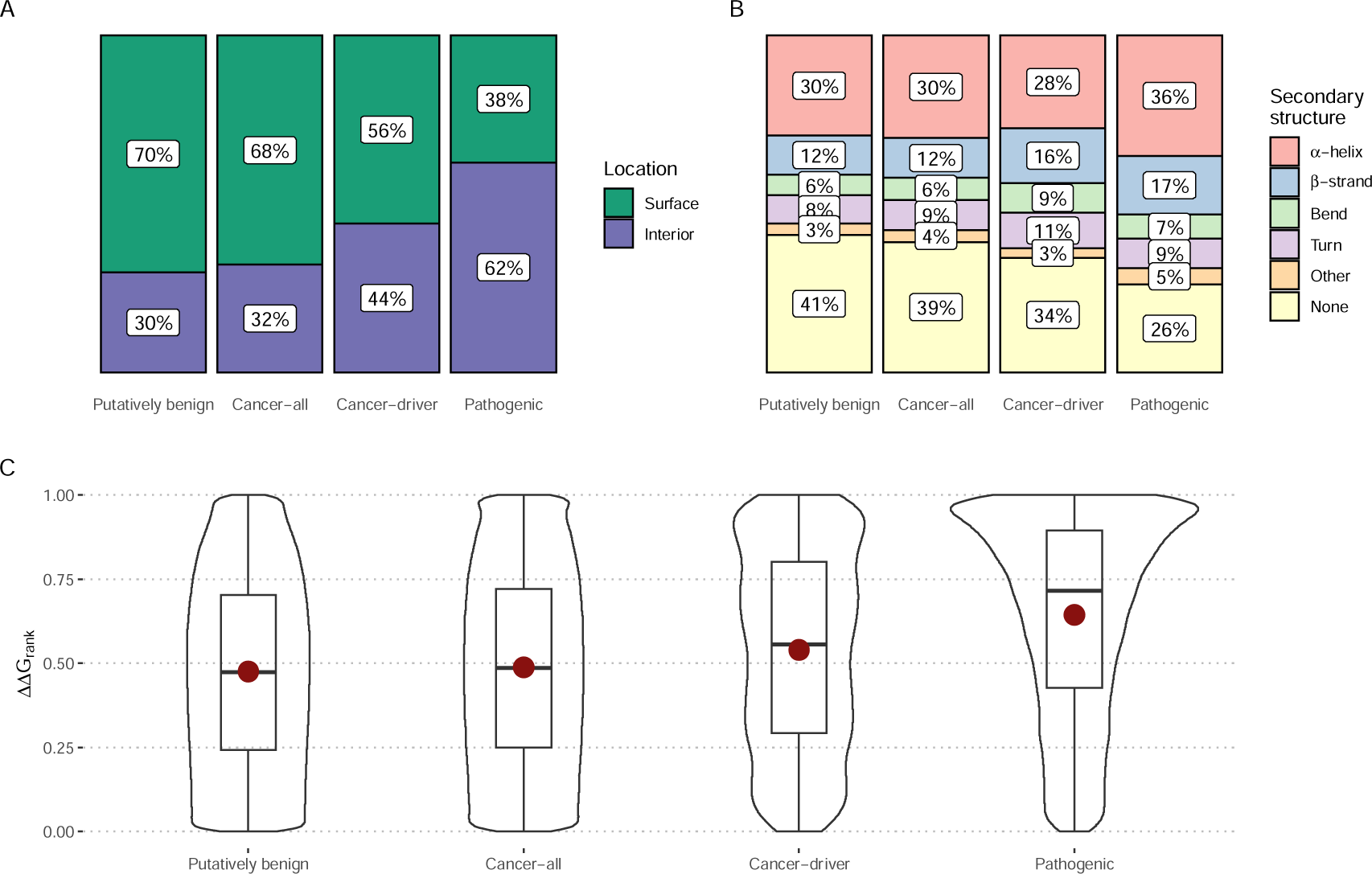
Protein structural properties of different types of missense mutations, based on AlphaFold models. Putatively benign mutations are those observed in the human population (gnomAD) without a reported disease association. Cancer-all are all cancer-associated mutations from the CMC. Cancer-driver mutations are the subset of cancer-associated mutations annotated for their direct role in cancer. Pathogenic mutations are those annotated as pathogenic or likely pathogenic in ClinVar. **(A)** Locations of mutations within protein structures in AlphaFold models, split in surface and interior as defined previously^74^. **(B)** Occurrence of mutations within different types of secondary structures. **(C)** Violin plot distributions of predicted structurally damaging effects, as measured by the ΔΔG_rank_ metric, whereby 0 represents the mildest possible single amino acid substitution in a protein, 1 represents the most damaging, and random mutations would be expected to have a mean of 0.5. The mean value of each distribution is represented with a red dot for the datasets. All iterations of group pairs proved to be highly significantly different (*p*-value < 3.3 x 10^-50^) according to Wilcoxon tests.

**Figure S2.**
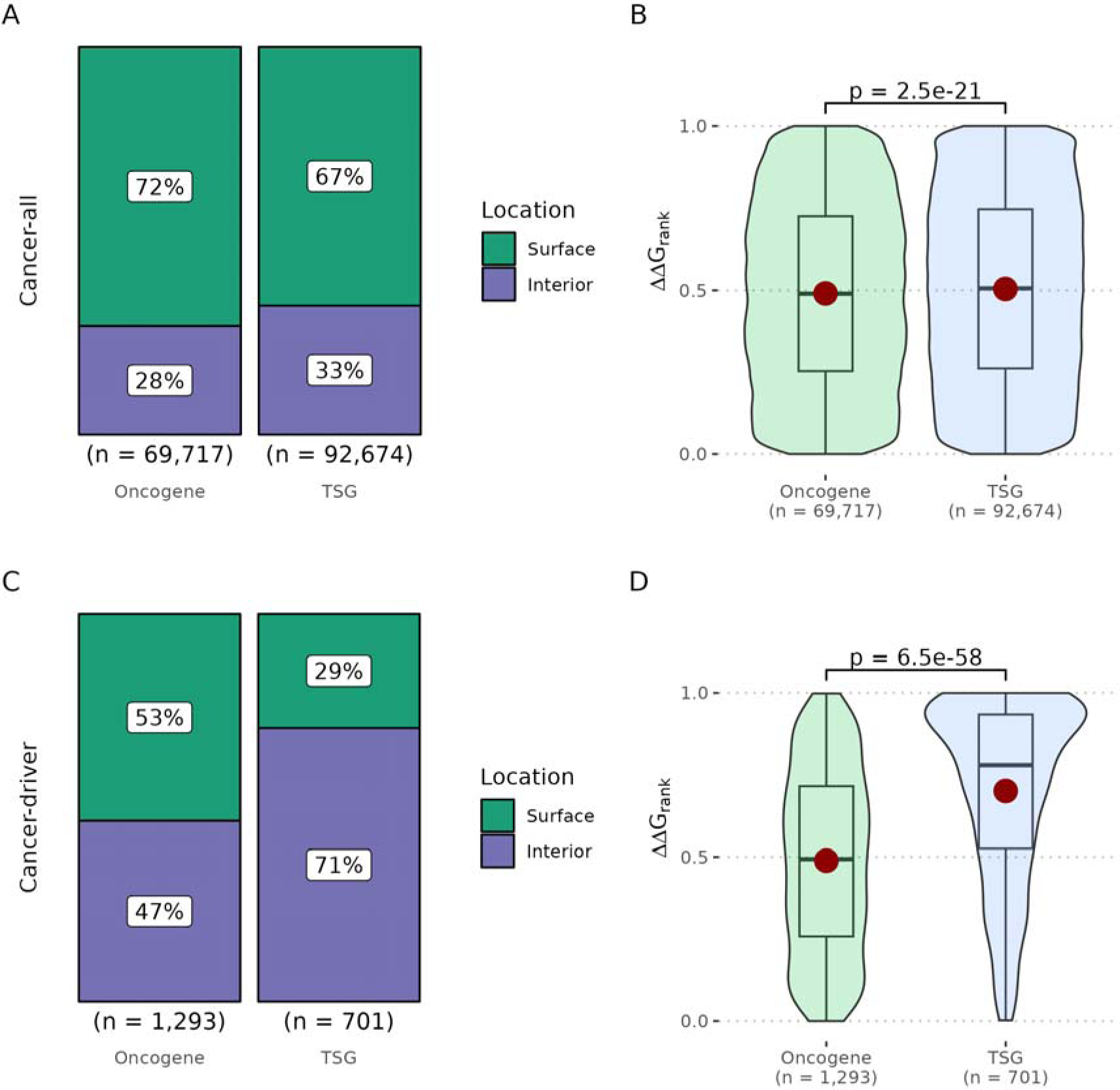
Protein structural properties of cancer-associated mutations in oncogenes and tumour suppressors, based on AlphaFold models. **(A)** Locations of all cancer-associated mutations within AlphaFold models (*cancer-all*) and **(B)** distributions of predicted structural damage, as represented by ΔΔG_rank_ values. **(C)** Locations of cancer-associated mutations with an annotated role in cancer (*cancer-driver*) and **(D)** distributions of ΔΔG_rank_ values. P-values were calculated using Wilcoxon tests.

**Figure S3.**
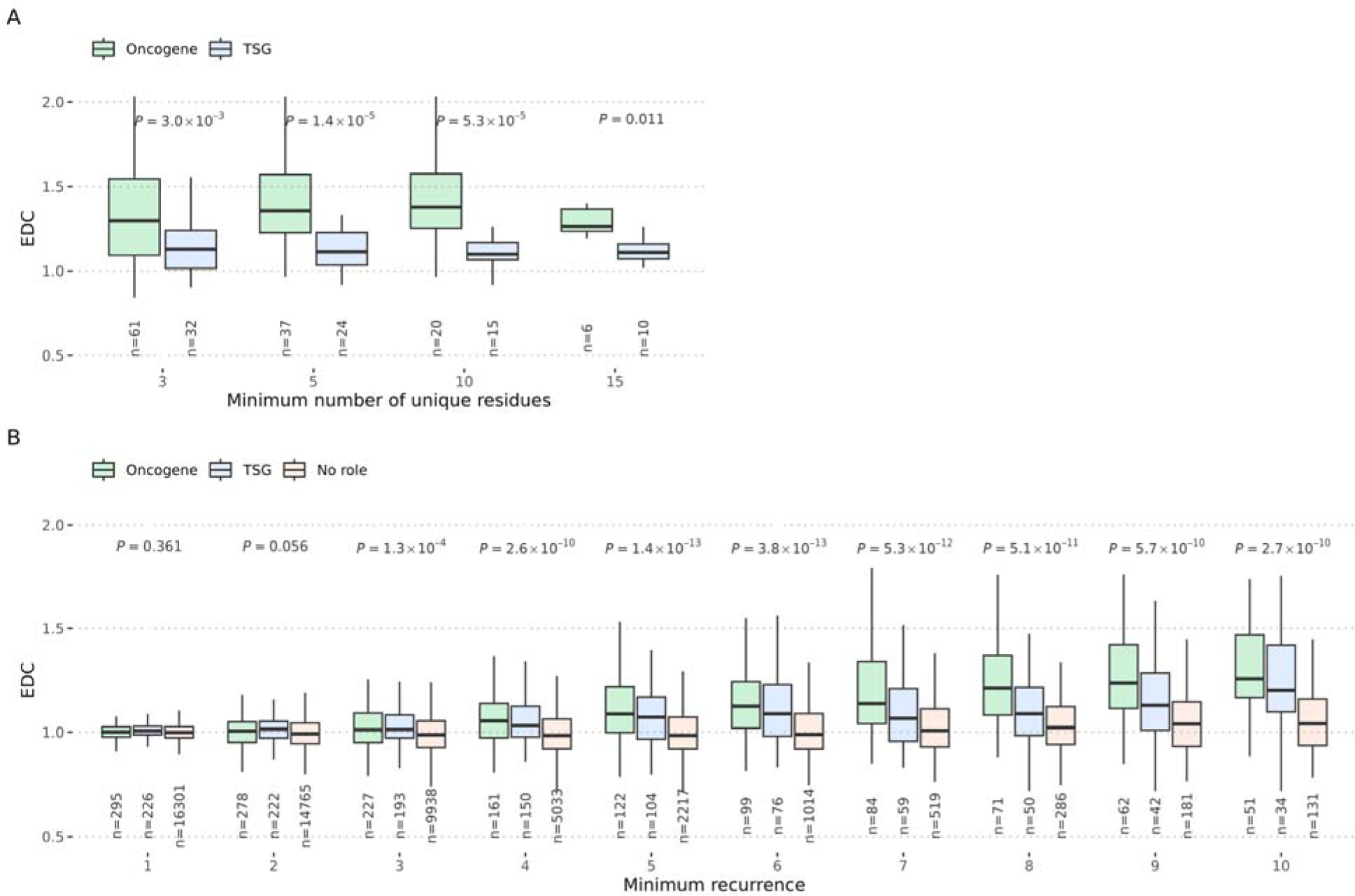
Thresholds for the number of unique and recurrent residues for calculating EDC. (**A**) The effect of the minimum number of unique residues on EDC values (at a minimum recurrence of 7). P-values were calculated with the Wilcoxon rank-sum test. (**B**) The effect of minimum recurrence (number of times a residue has been observed to be mutated in cancer samples) on EDC values (at a minimum number of unique residues >=5). P-values are given for comparisons between “Oncogene” and “No role” and were calculated with the Wilcoxon rank-sum test. Sample sizes at the bottom represent the number of genes in each group.

**Figure S4.**
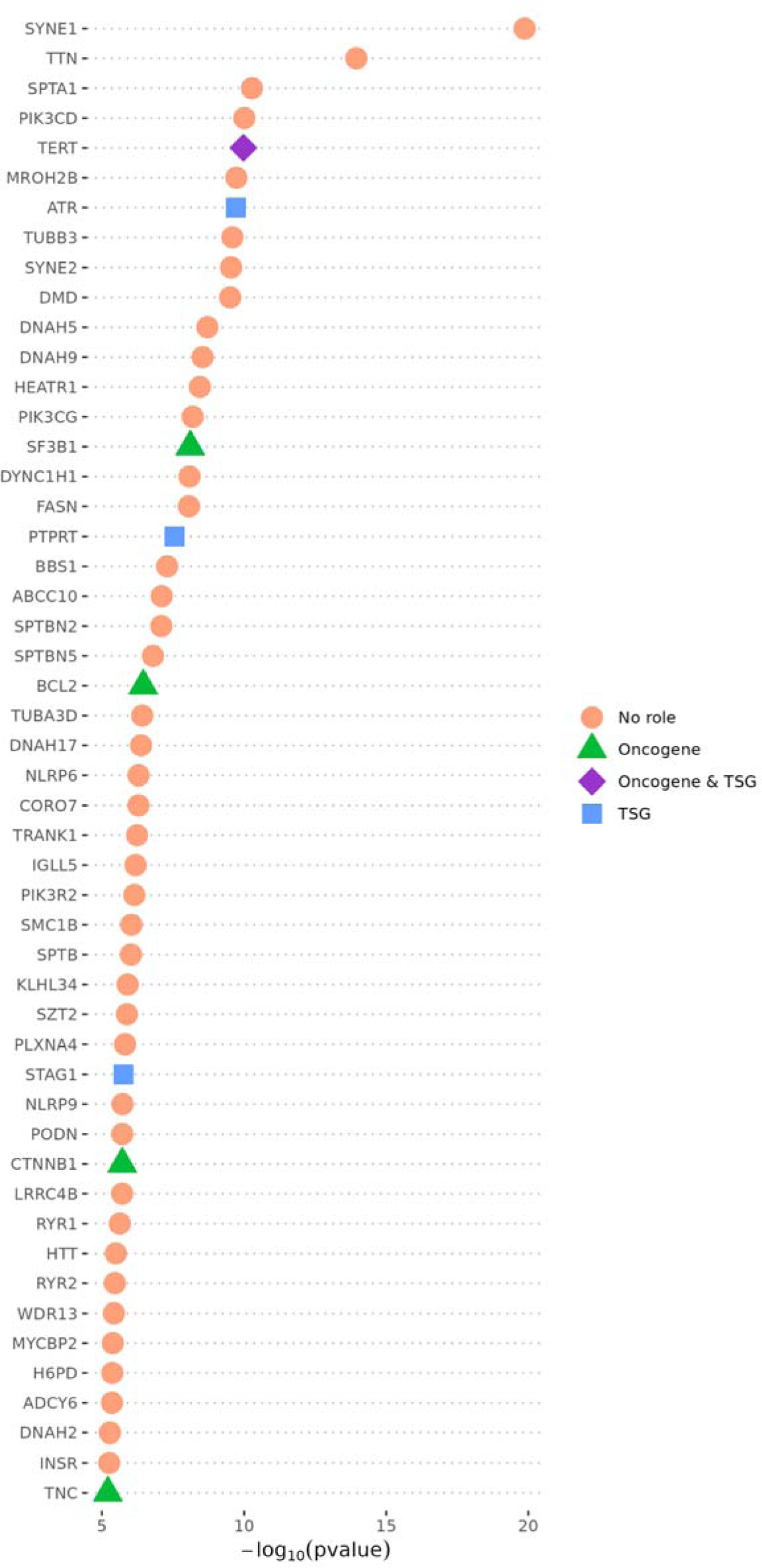
Top 50 human protein-coding genes enriched in structurally mild mutations. These are proteins with negative ΔΔG_rank_ difference values (*i.e.* those on the left side of the volcano plots in Fig. 3), for which the average observed missense mutation in the *cancer-all* dataset is less destabilising than the average unobserved but possible mutation. *P*-values from both the PDB and AlphaFold analyses in Fig. 3 are included, with the most significant value from either analysis selected for each protein to be used in this ranking.

